## Supplementary Materials and Methods for "Artificially inserted G-quadruplex DNA secondary structures induce long-distance chromatin activation"

Genomic Insertions using CRISPR-Cas9 genome editing

G4 array insert (*hTERT* promoter) (GC Percent- 76.73)

CCAGGCC**GGG**CT**CCC**AGTGGATtcgc**ggg**cacagacg**ccc**aggaccgcgctt**ccc**acgtggcgga**ggg**act**gggg**a**cccggg**ca**ccc**gtcctg**cccc**ttcaccttccagctccgcctcctccgcgcgga**cccc**g**cccc**gt**ccc**ga**cccc**t**cccggg**t**cccc**gg**ccc**ag**ccccc**tcc**gggccc**t**ccc**ag**cccc**t**cccc**ttcctttccgcgg**cccc**g**ccc**tctcctcgcggcgcgagtttcaggcagcgctgcgTCCTGCTGCGCACGT**GGG**AAGCC

G4 (G_3_L_1-20_ without bulge) mutated insert (GC Percent- 72.4)

CCAGGCC**GGG**CT**CCC**AGTGGATtcgc**gTg**cacagacg**ccc**aggaccgcgctt**ccc**acgtggcgga**gTg**act**ggTg**a**cccggg**ca**cAc**gtcctg**cccc**ttcaccttccagctccgcctcctccgcgcgga**cccc**g**cccc**gt**ccc**ga**cAcc**t**cAcgTg**t**cAcc**gg**cAc**ag**ccAcc**tcc**gggccc**t**ccc**ag**cAcc**t**ccAc**ttcctttccgcgg**cccc**g**ccc**tctcctcgcggcgcgagtttcaggcagcgctgcgTCCTGCTGCGCACGT**GGG**AAGCC

79M Left Homology Arm Primer for TERT G4 insert-

ATCCTAAGGAGATATAACTTCAAATATCCACCTCCAGGCCGGGCTCCCAGTGGAT

79M Right Homology Arm Primer for TERT G4 insert-

ATATCACAGGGTGTACACTATGTATGTACATCCGGCTTCCCACGTGCGCAGCAGGA

10M Left Homology Arm Primer for TERT G4 insert-

ACCTTGGCAGTCATATATTTTATCAATCCCTTGCCAGGCCGGGCTCCCAGTGGAT

10M Right Homology Arm Primer for TERT G4 insert-

GAATTAAATCTTCAAAATCCTTCCCTGAATCATGGCTTCCCACGTGCGCAGCAGGA

Homology arm

79M Adjacent primer F- GTACAGCCTACCTATGTGTACAAC

79M Adjacent primer R- CATGTGATATTAGAAGCAATAGCTC

10M Adjacent primer F- ATGTTCCATGAGTGGACCGC

10M Adjacent primer R- TGCACATTGCCTATACCAGGG

ChIP Primers

79M G4 array/mutated insert

R1 F- AAGTCGTAATTTTCTAGATGTTGCTT

R1 R- GCCTGGAGGTGGATATTTGA

R2 F- AAGTCGTAATTTTCTAGATGTTGCTT

R2 R- AAGGTGAAGGGGCAGGAC

R3 F- GGAAGCCGGATGTACATACATAGT

R3 R- TGTACACCCATCGTGATAAAAGA

R4 F- TTCTTTTATCACGATGGGTGTA

R4 R- ACACCATATGTGTTTTAATTAGAAGTG

10M G4 array/mutated insert

R1 F- ATAATGTTCCATGAGTGGACCGCA

R1 R- GGATGTGCAGTTGAATGCAGAAGGG

R2 F- TCCTCACCCTTCTGCATTCAACTG

R2 R- GAGGCGGAGCTGGAAGGTG

R3 F- CGGCGCGAGTTTCAGGCAG

R3 R- AAGCTCCATTTCAGGAACCGAACA

R4 F- ACGTGGGAAGCCATGATTCAG

R4 R- AGCACAGTCCCTCACATCACA

Promoters of the Surrounding Genes

PAWR F- CTCGTCGTATCGCCTGCT

PAWR R- CTCACACCCGGTCCTTTCC

PPP1R12A F- GACAGACCTGGAAAGGAGGA

PPP1R12A R- CATTCAAATAACGCGGCCAA

SYT1 F- GGGATGCTCTAGACCGAGTA

SYT1 R- CACCCTACACACACCCCTTC

NAV3 F- ACTGTTGTGGTCTTTTGGCT

NAV3 R- GGTAACTGCTACAAAGCTCCT

OSBPL8 F- GGCGTCCAAAGCTAACACTA

OSBPL8 R- AATGTGAAGGATCTTCGCGG

METTL25 F- CAGCCGACTAACTGCGTC

METTL25 R- AGGCTCAGTAGAGCTCCGA

ATXN7L3B F- GTTTGGCGGTGAGTCCTG

ATXN7L3B R- AGAGGGCGCTCGTTTTACT

SLC6A15 F- CAGCTTCTTGCCTAGGTTGG

SLC6A15 R- CAGCCTACCCTTCTCCTCTG

Gene (mRNA) expression

PAWR F- TGCCGCAGAGTGCTTAGATG

PAWR R- CCTGTAGCAGATAGGAACTGCC

PPP1R12A F- AGTTAATCGGCAAGGGGTTGA

PPP1R12A R- ATGACCACTATTTAGCCACTGC

SYT1 F- GCTGCTGGTAGGGATCATTCA

SYT1 R- GTTTTTCGGTGGACTTTTGTCTC

NAV3 F- GGATGTCCTAGAAGTCAGTCTCA

NAV3 R- GCTGCTTGTAGCGAGATAAACT

OSBPL8 F- TCCTCATAGCCAGGGTTTTGA

OSBPL8 R- AGGATCTGTGATTGTACTGAGCA

METTL25 F- TTGCTCTGGCTGCGAAATACT

METTL25 R- AACCCAAGTCAATCACCTGCT

ATXN7L3B F- GTGTGGCTACTTCTACCTGGA

ATXN7L3B R- GGCACGCTCCTTTGTCTTC

SLC6A15 F- AACTGGGCGGGATTGGATTTG

SLC6A15 R- GGTGGCAGAACTTTGTTCACATT

KLRC2 F- GCCAGCATTTTACCTTCCTCA

KLRC2 R- CACTGGGCTGATTTAAGTCGAT

KLRC3 F- GATTGGTGTGTTTCGTAACAGC

KLRC3 R- TCTGATGCACTGCAAGCTCAA

KLRC1 F- AGCTCCATTTTAGCAACTGAACA

KLRC1 R- CAACTATCGTTACCACAGAGGC

KLRK1 F- TTTTTCAACACGATGGCAAAAGC

KLRK1 R- GGGCCACAGTAACTTTCGGT

ETV6 F- ATCAACCTCTCTCATCGGGAA

ETV6 R- CAGTCTGCTATTCTCCCAATGG

NTF3 F- CCGTGGCATCCAAGGTAACAA

NTF3 R- GCAGTTCGGTGTCCATTGC

PTPRO F- ATGACTTCAGCCGTGTGAGAT

PTPRO R- GGGTGGCAATATACTCCTGGG

Chromosome Conformation Capture

PAWR + Insert F- CTGTAGTTTGCTGCCTAGATTCA

PAWR + Insert R- TTGTGGGAAACGTATGGCG

TaqMan Probe- AAGCTTTTTTAGAAGCAGAGTGACATGAC

PPP1R12A + Insert F- AGATTCAAGTCATGTCACTCTGC

PPP1R12A + Insert R- CTTACCATGGAAGTTTAGGGGTC

TaqMan Probe- CCTGACCCAGAACAGGTGCATAGT

SYT1 + Insert F- ATTCAAGTCATGTCACTCTGCTT

SYT1 + Insert R- GGAGACACAATCAAGGGCG

TaqMan Probe- TCTCCTGTGCATACTGCTTGCTGA

NAV3 + Insert F- AGTTTGCTGCCTAGATTCAAGT

NAV3 + Insert R- GCTATGACTTGGAGACACTGA

TaqMan Probe- GATGAAGCTAGAAAGGGGGAAAGAATGTATG
